# Modern microbialites harbor an undescribed diversity of chromerid algae

**DOI:** 10.1101/2025.07.02.662897

**Authors:** Anthony M. Bonacolta, Patrick J. Keeling

## Abstract

Chromerid algae and the heterotrophic colpodellids together make up the chrompodellids, which are the closest known relatives to apicomplexan parasites [1]. As apicomplexans are prolific parasites of animals, including humans, their adaptation to parasitism from algal ancestors has garnered significant interest, and chromerids were instrumental in elucidating that evolution [2-5]. But the chromerids were also the first new algal group discovered in 100 years [2], and because so much attention has focused on comparisons with apicomplexans, the ecology of chromerid algae has remained surprisingly mysterious. They have predominately been considered to live only in association with corals, initially as intracellular coral symbionts. To test for a wider distribution of chromerids in nature, we have used plastid metagenomic binning combined with re-analysis of 18S rRNA metabarcoding data, which both support an expanded biogeographic range and ecological niche of chromerid algae by showing they are consistent associates of modern microbialites across the globe. Specifically, a broad and undescribed diversity of chromerid lineages is found in marine and also freshwater microbialites. This includes the recovery of a near-complete *V. brassicaformis* plastid genome and a second incomplete plastid genome from a new chromerid lineage from Highborne Cay thrombolites. This is the first concrete evidence that this under-sampled algal lineage lives outside some association with coral, including freshwater habitats, and sheds new light on the diversity and ecological range of the most recently discovered algal lineage.

## Main

Since their discovery almost two decades ago, there are still only two described chromerid algae, *Chromera velia* [6] and *Vitrella brassicaformis* [7]. While these two species were both initially isolated from coral in Australia, significant divergence in their morphologies, life histories, and evolution have always been apparent [7]. Nuclear gene phylogenies also support this by showing they are distantly related within the larger chrompodellid lineage: *V. brassicaformis* branches near the base of the chrompodellids with the mollusk parasite *Piridium sociable*, whereas *C. velia* branches with free-living predators *Colpodella, Voromonas*, and *Alphamonas* [1, 2]. Their plastid genomes are similarly divergent, with *C. velia* bearing a linear, 120 kb genome, while *V. brassicaformis* possesses a circular gene-rich 85 kb genome [8]. Their ecological roles are also seemingly divergent, but here we have much less data to go on: despite being the first new algal group to be discovered in almost 100 years, we know almost nothing about their role in natural environments and critical questions, especially their association with coral, are still debated. After first being presumed to be in an intracellular photosymbiotic association with coral [6, 9], the transcriptomic response of coral larvae exposed to *C. velia* was shown to more resemble parasite invasion, and markedly low colonization levels were observed compared to Symbiodiniaceae [10, 11]. Closer analyses of environmental data also suggested most chromerid sequences are more consistent with growth on coral surfaces, as opposed to being directly associated with coral tissue [12, 13]. The identification of chromerid and other apicomplexan-related lineages (ARLs) in environments outside coral reefs further supports a broader ecological range for these algae [12, 14]. For instance, *Vitrella*-related 18S and plastid 16S rRNA gene sequences (classified as ARL-I), are mostly found on coral reefs, but are also present in a range of calcium carbonate-rich environments like reef sediments and even microbialites [1, 12, 14].

Microbialites are microbially-induced formations of trapped, bound, or precipitated sediment that exhibit a range of mineralogies, although calcium carbonate is the most widespread [15]. Ancient microbialites offered some of the first evidence for carbonate-based reef communities, and modern microbalites offer fascinating insights into early evolution [16–18].

The potential association between chromerid algae and microbialites is tantalizing given this evolutionary context and compositional similarity between coral and microbialite reef systems. To further study this link and to gather additional information on the biogeography of chromerid algae, we employed new genomic binning approaches to shotgun metagenomic datasets, and re-analyzed existing 18S rRNA gene datasets from microbialite environments, most of which were not previously reported to include chromerids.

Metagenomic reads from Highborne Cay (Bahamas) thrombolites were downloaded from NCBI SRA (BioProject: PRJNA1055967), trimmed using Trimmomatic v0.39 [19], then assembled using MegaHit v1.2.9 [20]. *Vitrella-*related and *Chromera*-related 18S rRNA genes were identified and extracted using Blastn [21], then aligned with other chrompodellid and apicomplexan rRNA operons (18S + 28S rRNA genes) using MAFFT v7 [22]. A maximum-likelihood backbone tree was constructed using IQ-tree v2.1.0 with the GTR+F+R5 model (as selected by ModelFinder) [23]. Amplicon sequence variants (ASVs) classified as chromerids were extracted from 18S rRNA gene datasets originating from microbialites found in Pavilion Lake (Canada), Kelly Lake (Canada) [18], Lake Alichichica (Mexico) [24], Highborne Cay (Bahamas), and Shark Bay (Australia; Figure 1) [25]. These microbialite are distributed across the globe and range in salinity from freshwater to hypersaline, representing both geographically and chemically distinct environments. These ASVs were placed using RAxML’s evolutionary placement algorithm [26] onto the well-supported backbone tree of the apicomplexan + chrompodellid rRNA operons to assess chromerid biodiversity across global microbialites (Figure 1). The full-length *Vitrella-*related 18S rRNA gene from Highborne Cay branched within a well-supported (100% bootstrap) clade with other *V. brassicaformis* sequences recovered from Australian corals and the Arabian Sea coral reef sediments. These sequence and other environmental sequences in this clade showed minimal divergence, suggesting they all likely belong to the same species. In contrast, however, other *Vitrella*-related ASVs from marine microbialites all branch outside this clade, and likely represent a distinct species of *Vitrella*. The *Chromera*-related 18S rRNA gene recovered from Highborne Cay branches sister to *C. velia* but likely represents a different species or possibly even genus based on phylogenetic divergence (Figure 1). Strikingly, we found freshwater and marine chromerid ASVs from microbialites cluster with this sequence as well, distinct from *C. velia* (Figure 1). Biological support for the freshwater ASV (ASV_3680) is strong considering it is found in multiple samples across two different locations [18]. Its presence amongst marine-associated microbialite ASVs further supports this clade as a likely unexplored, yet globally distributed *Chromera*-related lineage associated specifically with microbialite environments. The hypersalinity of shark bay microbialites inhibits coral reef growth [27], thus providing additional evidence for a microbialite-specific association, as opposed to contamination from nearby coral reef environments. While *Vitrella* spp. have been noted in microbialite environments previously [14], they were never observed to be abundant or globally distributed, and this is the first definitive evidence of *Chromera*-related sequences recovered outside of a coral reef habitat.

**Figure 1.**
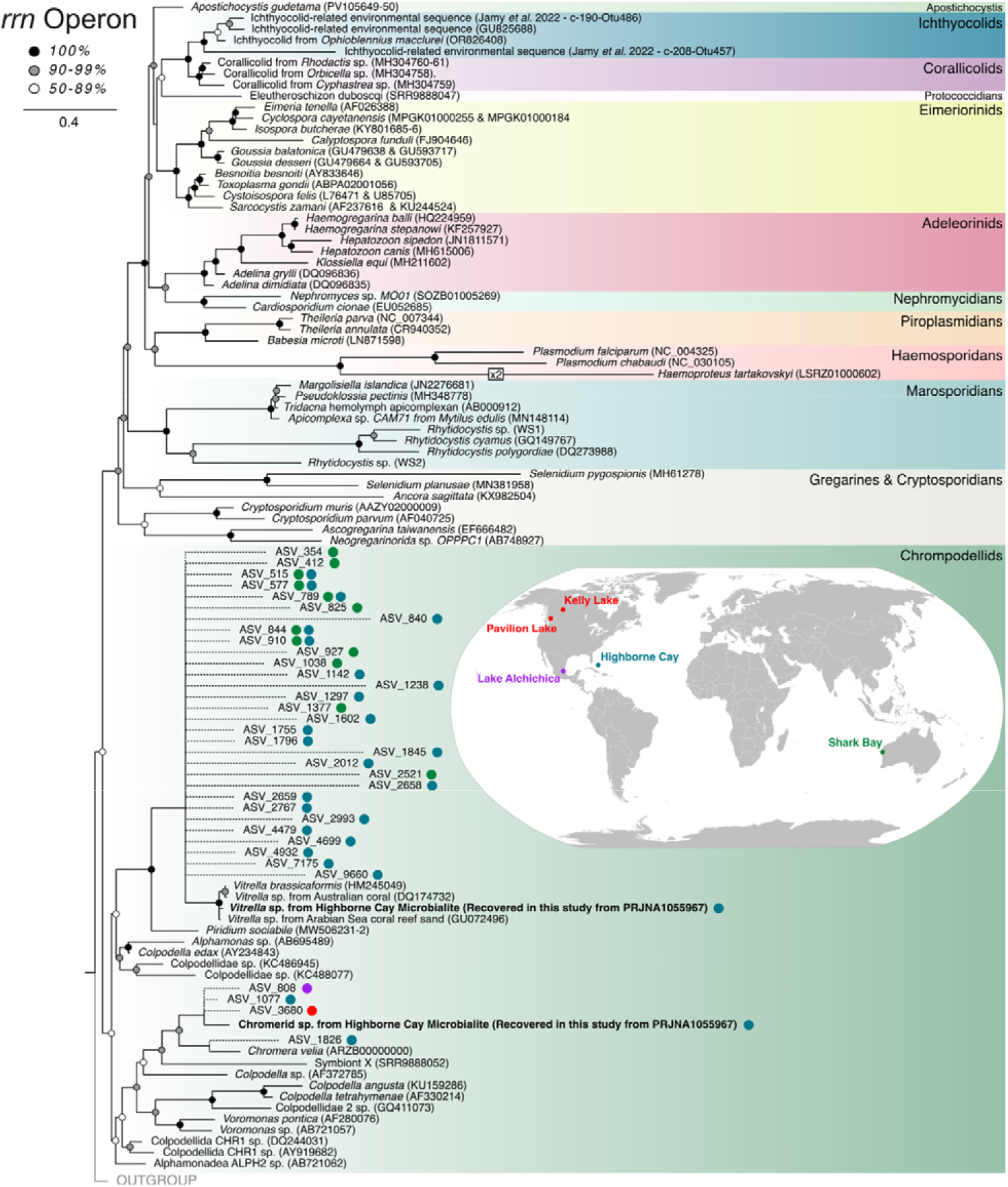
Maximum-likelihood tree of apicomplexans and chrompodellids based on the nuclear rRNA operon with EPA-placed ASVs recovered from microbialite 18S rRNA gene metabarcoding studies. World map (inset) showing locations of 18S rRNA gene metabarcoding datasets analyzed in this study. The colored dots next to each ASV represents the origin of each sequence. Note that ASV_3680 (red dot) was found in both Canadian microbialite locations.

Using Metabat v2.18 [28] optimized for the retrieval of plastid metagenome assembled genomes (ptMAGs; -s 30000 --minContig 1500), we recovered two ptMAGs corresponding to chromerid plastids from Highborne Cay thrombolites. Extracting the plastid *rrn* operon (16S + 23s rRNA genes) from both ptMAGs revealed similar patterns to the 18S rRNA genes we retrieved, where one ptMAG *rrn* operon corresponds to *V. brassicaformis* and the other corresponds to a chromerid closely related to but distinct from *C. veli*a (Figure 2A). The *Chromera*-related ptMAG was partial, at 15 kb, but nevertheless contained *psbE, SecA, psaC*, and *atpH* (Figure 2B), the phylogenies of which confirmed each to be closely related to *C. velia* and distinct from *Vitrella*. The presence of photosystems within this ptMAG also confirms the organism is photosynthetic, altogether demonstrating the presence of uncultured, photosynthetic *Chromera*-related algae in modern microbialites. The ptMAG corresponding to *V. brassicaformis* is ∼82 kb and contained a majority of *V. brassicaformis* plastid genes (which were also confirmed by phylogenetic analyses to be closely related to *V. brassicaformis* homologues), however it could not be assembled into a single contig or circularized (Figure 2C). A phylogenetic tree based on 31 plastid genes confirmed that this ptMAG is very closely related to *V. brassicaformis* (Figure 2C), confirming that photosynthetic algae from this lineage live in calcium carbonate habitats beyond coral reefs.

**Figure 2.**
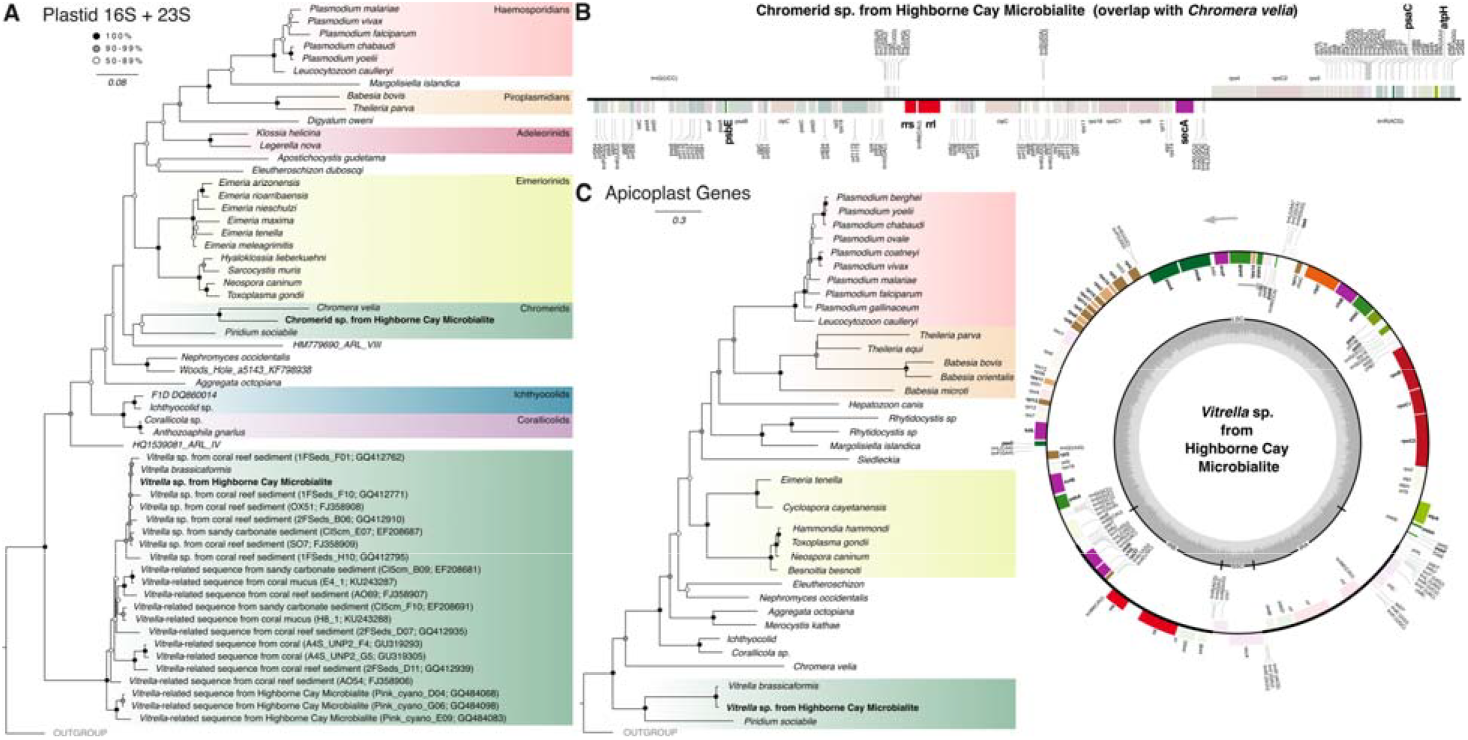
(A) Maximum-likelihood tree of apicomplexans and apicomplexan-related lineages based on the plastid *rrn* operon (16S + 23S rRNA genes). (B) Novel *C. vela*-related plastid genome assembly recovered from a Highborne Cay microbialite MAG (metagenome assembled genome). Recovered genes (bold, fully coloured) are overlayed on the *Chromera velia* plastid genome. (C) A *V. brassicaformis*-related plastid genome MAG from Highborne Cay microbialites. On the right is recovered *Vitrella brassicaformis* plastid genes (bold, fully coloured) are overlayed over the *V. brassicaformis* plastid genome. On the left is a maximum-likelihood tree of apicomplexans and apicomplexan-related lineages based on 31 plastid-encoded genes, including genes recovered from the *V. brassicaformis*-related plastid genome. Plastid diagrams were generated using OGdraw v1.3.1 [32]

Overall, these analysis show that *V. brassicaformis* and a novel, photosynthetic chromerid alga live within modern microbialites, expanding the known distribution of both subgroups of chromerid algae beyond coral reef environments. Both the current data and previous analyses of environmental ASV data show that chromerids are not normally highly abundant in most environments (with rare exceptions [29]), but their consistent presence in similar but geographically distant environments is evidence of an important ecological niche fulfilled by these algal lineages. *Ostreobium qurkettii*, another coral-associated epiphytic alga [30], is also found consistently within marine microbialites [18], which may offer clues as to the ecological function of chromerids in this environment. Twenty years of research on these algae has yielded significant insights, but mostly related to the evolution of apicomplexans and their plastids. Only two species of chromerids have been cultured or described thus far, none in the last decade, and very little insight into their ecological roles in nature have been made beyond their initial discovery. We suggest that the cryptic chromerid diversity of modern microbialites should be prioritized in future culturing efforts. In culture, *C. velia* tends to form a brownish, endophytic layer on the cultivation flask [31] and the morphological form of *C. velia* is influenced by salinity [9], suggesting that the undescribed chromerids of hypersaline microbialites (i.e. Shark Bay, Australia) may be observed as non-motile occoids. From what we know about the drastic differences in morphology and genome evolution of the two cultured chromerids, a new culture of the novel *Chromera*-related algae would further expand the diversity of various biological characteristics of the group still more, and a more detailed appraisal of their ecological and physical position within the microbialite community should shed light on these characteristics of the group more broadly in more complex reef communities.

## Data availability

The raw data used for this project was retrieved from the NCBI SRA database. Partial plastid assemblies, SGTs, and raw tree files can be found on GitHub at: https://github.com/Abonacolta/microbialite_chromerids. Recovered 18S rRNA genes have been deposited onto NCBI GenBank under the following accession numbers: PV865597& PV865598.

## Acknowledgments

This research was enabled in part by support provided by the Digital Research Alliance of Canada (alliancecan.ca). This project was supported by a grant from the Gordon and Betty Moore Foundation (https://doi.org/10.37807/GBMF9201).

